# Low Replication Capacity Virus is Preferentially Transmitted in Mother-to-Child-Transmission but not in Adult-to-Adult-Transmission of HIV-1

**DOI:** 10.1101/510776

**Authors:** Emily Adland, Louisa Iselin, Francesca Roxburgh, Jane Millar, Anna Csala, Vanessa Naidoo, Christie Noble, Jake Thomas, Alessia De’Felice, Ago Szabo, Nick Grayson, Amber Moodley, Krista Dong, Bruce Walker, Thumbi Ndung’u, Samantha McInally, Eric Hunter, Philip Goulder

## Abstract

Previous studies of the transmitted/founder virus compared to viral quasispecies in the donor have yielded conflicting results. In heterosexual adult-to-adult transmission (ATAT), the viral replicative capacity (VRC) of transmitted virus is reportedly either similar to, or somewhat higher than, that of donor virus, whilst transmitted virus in mother-to-child transmission (MTCT) has a significantly lower VRC than that of maternal virus. These discrepancies may be explained by the different methodologies used in these studies, or they may reflect true differences in the transmission bottleneck. To resolve this question, we here use the same methodology to compare transmitted versus donor virus in MTCT and ATAT. We show that, in a South African mother-child cohort, infant virus samples obtained at 1-2 days after birth had VRC significantly lower than in the mothers (p=0.0003). By contrast, in Zambian ATAT transmission pairs, VRC of transmitted virus was similar to or somewhat higher than donor virus (p=ns). The VRC of virus transmitted to the recipient, compared to that in the donor, was significantly lower in MTCT versus heterosexual ATAT (p=0.01). These studies demonstrate that fundamental differences exist between the viruses transmitted via the MTCT and ATAT bottlenecks that are not explained by methodological factors. This result is of importance since transmission of low replicative capacity virus results in low immune activation and a small viral reservoir, and therefore the preferential transmission of low fitness viruses in MTCT might be expected to increase cure potential in *in utero* infected infants and children.

**IMPORTANCE:** Understanding the factors determining which viruses are preferentially transmitted in HIV infection is critical to the development of new, effective strategies to prevent transmission. Despite this, much remains unknown in this respect, both with regard to adult-to-adult transmission (ATAT) but especially with respect to mother-to-child transmission (MTCT). The finding here that fundamental differences exist in the genetic bottleneck of HIV transmission between heterosexual ATAT and MTCT is an important initial step to help define the viral mechanisms contributing to transmission in each case. In addition, we show that viruses of low viral replicative capacity are preferentially transmitted in MTCT. This suggests the possibility that a window of opportunity exists following *in utero* infection in which early anti-viral intervention not only reduces the size and diversity of the viral reservoir, but additionally maintains a reservoir comprising low viral replicative capacity HIV. Low replicative capacity of transmitted virus has previously been shown to result in low immune activation and low proviral DNA load in central memory cells, factors likely to be directly relevant to increasing cure potential in HIV-infected infants and children.

## INTRODUCTION

Analysis of the differences between HIV infection in children and adults has facilitated understanding concepts such as HIV pathogenesis and cure (1–6). In addition, defining the distinct mechanisms of HIV transmission in adult-to-adult (ATAT) versus mother-to-child transmission (MTCT) is directly relevant to the development of effective strategies to prevent HIV infection in adults and children. Reportedly, one substantial difference between ATAT and MTCT appears to be in the replicative capacity of the transmitted virus. In MTCT, the transmitted/founder virus has a significantly lower replicative capacity than virus in the mother (7, 8). By contrast, in adult-to-adult heterosexual transmission, the replicative capacity of transmitted/founder virus has been reported as similar to, or somewhat higher than, virus in the donor (9, 10). In part, the differences between these studies may be explained by the distinct methodologies that have been used. For this reason, we set out here to determine whether, using the same methodology, the MTCT bottleneck selects for lower replication capacity virus than the ATAT bottleneck in heterosexual transmission.

In order to investigate potential differences in virus transmission comparing MTCT and ATAT, we measured Gag-protease-driven viral replication capacity (7, 11) in a cohort of *in utero* infected infants and their mothers in KwaZulu-Natal, South Africa, from a timepoint a median of 36hrs after birth, and first compared these to the ‘FRESH’ cohort of acutely infected women also from KwaZulu-Natal (12) in which the donor was not known. We then evaluated the reduction in replication capacity of transmitted/founder virus versus donor virus in the mother-child transmission pairs compared with a cohort of adult heterosexual transmission pairs from Zambia. Finally, to determine whether this method of establishing viral replicative capacity (VRC) might artefactually inflate the difference between donor and recipient both in MTCT and ATAT, in each of these two cohorts we assessed the ability of the Gag-Protease-driven viral replication assay, that uses population Gag-Pro sequences, to represent the replication capacity of individual viral clones generated from the same individual.

## MATERIALS AND METHODS

### Patients and samples

We studied viral replicative capacity in a total of 86 mother infant transmission pairs, 15 acutely infected adult females and 9 adult transmission pairs, all with HIV-1 C-clade infection.

#### Ucwaningo Lwabantwana Study, South Africa

58 mother infant pairs were recruited between 2015 and 2017 from 4 study sites in South Africa, Edendale Hospital, Pietermaritzburg; Stanger Hospital, Kwa-Dukuza; Queen Nandi Hospital, Empangeni, and Mahatma Gandhi Memorial Hospital, Durban. The mean absolute CD4 count of the mothers was 456/mm^3^ (interquartile range (IQR) 255-566/mm^3^) and the median viral load 8,000 copies/ml (IQR <20-770,000). The median age of the infants was 1.54 days (IQR 0.89-9 days), mean CD4% 46% (IQR 22-51%), mean absolute CD4 count 1872/ mm^3^ (IQR 1099-2446/ mm^3^) and median viral load 18,000 copies/ml (IQR 2300-99,500).

#### PEHSS Study, South Africa

28 *in utero* infected mother-infant pairs were recruited from St Mary’s Hospital, Mariannhill, Durban and Prince Mshiyeni Hospital, Umlazi, Durban South Africa between 2002 and 2005 (13, 14). The mean absolute CD4 count of the mothers was 375/mm^3^ (IQR 211-483/mm^3^) and the median viral load 49,750 copies/ml (IQR 11,800-169,250). The mean age of the infants was 6 weeks (range 4-8 weeks), mean CD4% 33% (IQR 26-40%), mean absolute CD4 count 1581/ mm^3^ (IQR 1081-2108/ mm^3^) and median viral load 91,550 copies/ml (IQR 24,675-1,198,000).

#### Females Rising through Education, Support, and Health (FRESH) Study, South Africa

Between 2012 and 2015 HIV-negative women aged 18-23 years were recruited for the FRESH study in Umlazi, South Africa (12, 15). Assessment of HIV status was made at study entry and twice per week during follow up with a rapid screening HIV RNA PCR assay followed by a confirmatory assay. For the purposes of this study we measured VRC in 15 acutely infected women, the mean absolute CD4 count of the women was 642/mm^3^ (IQR 390-903/mm^3^) and the median viral load was 128,500 copies/ml (IQR 6,250-380,000).

#### Zambia Emory HIV Research Project (ZEHRP), Zambia

The 9 transmission pairs selected for this study were enrolled in the heterosexual discordant couple cohort at the Zambia-Emory HIV Research Project in Lusaka, Zambia (16, 17). Each recipient of the transmission pair had plasma samples available within 32 days of the estimated date of infection (15-32 EDI) and showed previous evidence of a single-variant infection (infection was established by one transmitted-founder virus). In addition, the donors had samples available within 50 days of the estimated date of infection of the recipient (8-45 EDI). The current study comprised of 5 male-to-female and 4 female-to-male transmission pairs. The mean absolute CD4 count of the 9 donors was 500/mm^3^ (Range 115-1028/mm^3^) and the median viral load was 79,264 copies/ml (Range 2,620-365,000). The absolute CD4 count of the 9 recipients was 507/mm^3^ (Range 320-828/mm^3^) and the median viral load was 2,941,142 copies/ml (Range 78,600-24,993,584).

#### Viral load and absolute CD4 count measurement

Viral load measurement was undertaken using the BioMérieux NucliSens Version 2.0 Easy Q/Easy Mag (NucliSens v2.0) assay (dynamic range 20-10m) prior to 2010 and using the COBAS Ampliprep/COBAS TaqMan HIV-1 Test version 2.0 by Roche (CAP/CTM v2.0) (dynamic range 20-10m subsequent to 2010. CD4+ T cell accounts were measured by flow cytometry.

#### Consent

Informed consent was provided for participation of the subjects in each of the studies. Ethics approval was given by the Biomedical Research Ethics Committee, University of KwaZulu-Natal, Durban and the University of Zambia Research Ethics Committee and the Emory University Institutional Review Board.

### Viral RNA isolation and nested RT-PCR amplification of population *gag-protease* from plasma

Viral RNA was isolated from plasma by use of a QIAamp Viral RNA Mini Kit from Qiagen. The *gag-protease* region was amplified by reverse transcription (RT)-PCR from plasma HIV-1 RNA using Superscript III One-Step Reverse Transcriptase kit (Invitrogen) and the following *gag-protease* -specific primers: 5’ CAC TGC TTA AGC CTC AAT AAA GCT TGC C 3’ (HXB2 nucleotides 512 to 539) and 5’ TTT AAC CCT GCT GGG TGT GGT ATT CCT 3’ (nucleotides 2851 to 2825). Second round PCR was performed using 100-mer primers that completely matched the pNL4-3 sequence using Takara EX Taq DNA polymerase, Hot Start version (Takara Bio Inc., Shiga, Japan). One hundred microliters of reaction mixture was composed of 10ul of 10x EX Taq buffer, 4ul of deoxynucleoside triphosphate mix (2.5 mM each), 6ul of 10uM forward primer (GAC TCG GCT TGC TGA AGC GCG CAC GGC AAG AGG CGA GGG GCG GCG ACT GGT GAG TAC GCC AAA AAT TTT GAC TAG CGG AGG CTA GAA GGA GAG AGA TGG G, 695 to 794) and reverse primer (GGC CCA ATT TTT GAA ATT TTT CCT TCC TTT TCC ATT TCT GTA CAA ATT TCT ACT AAT GCT TTT ATT TTT TCT TCT GTC AAT GGC CAT TGT TTA ACT TTT G, 2646 to 2547), 0.5ul of enzyme, and 2ul of first round PCR product and DNase-RNase-free water. Thermal cycler conditions were as follows: 94°C for 2 min, followed by 40 cycles of 94°C for 30 s, 60°C for 30 s, and 72°C for 2 min and then followed by 7 min at 72°C. PCR products were purified using a QIAquick PCR purification kit (Qiagen, UK) according to manufacturer’s instructions.

### Single Genome Amplification

For single genome amplification of the gag-protease genes, RNA was endpoint diluted in 96-well plates such that fewer than 29 PCRs yielded an amplification product. According to a Poisson distribution, the cDNA dilution that yields PCR products in no more than 30% of wells contains one amplifiable cDNA template per positive PCR more than 80% of the time. RT-PCR and second round PCR were carried out as described above for the population gag-protease amplification. PCR products were visualized by agarose gel electrophoresis under UV light using a Gel Doc 2000 (Biorad). All products derived from cDNA dilutions yielding less than 30% PCR positivity were sequenced. For the 27 adult and infant samples for which the comparisons were made with population-derived VRC, the number of SGA clones analysed per subject was a median of 6 (IQR 5-7).

### Generation of recombinant Gag-Protease viruses

A deleted version of pNL4-3 was constructed (18) that lacks the entire Gag and Protease region (Stratagene Quick-Change XL kit) replacing this region with a BstE II (New England Biolabs) restriction site at the 5’ end of Gag and the 3’ end of protease (11). To generate recombinant viruses, 10ug of BstEII-linearized plasmid plus 50ul of the second-round amplicon (approximately 2.5ug) were mixed with 2 x 10^6^ cells of a Tat-driven green fluorescent protein (GFP) reporter T cell line (19) (GXR 25 cells) in 800ul of R10 medium (RPMI 1640 medium containing 10% fetal calf serum, 2 mM L-glutamine, 100 units/ml penicillin and 100ug/ml streptomycin) and transfected by electroporation using a Bio-Rad GenePulser II instrument (300 V and 500 uF). Following transfection, cells were rested for 45 min at room temperature, transferred to T25 culture flasks in 5ml warm R10 and fed with 5ml R10 on day 4. GFP expression was monitored by flow cytometry (LSR II; BD Biosciences), and once GFP expression reached >30% among viable cells supernatants containing the recombinant viruses were harvested and aliquots stored at −80°C.

### Replication Capacity Assays

The replication capacity of each chimera is determined by infection of GXR cells at a low multiplicity of infection (MOI) of 0.003 (7, 11). The mean slope of exponential growth from day 2 to day 7 was calculated using the semi log method in Excel. This was divided by the slope of growth of the wild-type NL4-3 control included in each assay to generate a normalized measure of replication capacity. Replication assays were performed in triplicate, and results were averaged. Importantly, all the VRC determinations described here were undertaken blinded to the identity of the study subject.

### Viral sequencing and phylogenetic analysis

Sequencing was undertaken using the Big Dye ready reaction terminator mix (V3) (Department of Zoology, University of Oxford). Sequence data were analysed using Sequencher v.4.8 (Gene Codes Corporation). Nucleotides for each gene were aligned manually in Se-Al v.2.0a11. Maximum-Likelihood phylogenetic trees were generated using PHYm131 (http://www.hiv.lanl.gov) and visualised using Figtree v.1.2.2 (http://tree.bio.ed.ac.uk/software/figtee/).

### Statistical Analysis

The VRCs between mother-child pairs, adult transmission pairs, and between SGA-derived and population-derived VRCs, were compared using the paired t test or, where data were non-normally distributed, using the Wilcoxon matched-pairs signed rank test. The differences in VRC between donor and recipient among MTCT pairs and heterosexual ATAT pairs was determined using the Mann Whitney test. The relationship between population-derived VRC and SGA-derived VRC were assessed using the Spearman’s correlation.

## RESULTS

### Lower viral replicative capacity in infants compared to the transmitting mothers

Chimeric viruses generated by co-transfection of the amplified patient-derived gag-protease genes with linearized pNL4-3Δgag-protease were used to infect a GFP reporter cell line as previously described (7, 11). Recombinant viruses were generated from mother-infant pairs from the PEHSS cohort (8), of whom 28 infants were *in utero* infected (mean infant age 6 weeks). We now also generated recombinant viruses from 58 mother-infant pairs from the UL cohort (mean infant age 37 hours), and 15 acutely infected adult females from the FRESH cohort (Table 1). Several previous studies adopting this methodology (7, 11, 18, 20, 21) have demonstrated 99-100% amino acid sequence identity between plasma virus and the Gag-Pro-chimeric viruses, after exclusion of codons containing mixed bases. Thus, the Gag-Pro-chimeric viruses that were generated are indeed representative of the plasma virus in each study subject.

**Table 1:** Clinical parameters of the 4 cohorts, PEHSS, UL, FRESH and ZEHRP, of HIV infected adults and infants used in this study. Clinical data shown for n=58 mother-infant transmission pairs from Ucwaningo Lwabantwana (UL) Study, South Africa, n=28 mother-infant transmission pairs from Paediatric Early HAART and Structured Interruption Study (PEHSS), South Africa, n=15 acutely infected female adults from the Females Rising through Education, Support, and Health (FRESH study), South Africa and n=9 acutely infected adult-to-adult transmission pairs from the Zambia Emory HIV Research Project (ZEHRP), Zambia. (ACC) absolute CD4 count; (VL) viral load expressed as copies/ml plasma. Values shown are the mean of the group and IQR in brackets.

| Cohort | MTCT/ATAT | n | Donor Sex | Donor Age | Donor CD4 | Donor viral load | Recipient Sex | Recipient Age | Recipient CD4 | Recipient viral load |
| --- | --- | --- | --- | --- | --- | --- | --- | --- | --- | --- |
|  |  |  |  |  | /mm<sup>3</sup> | HIV RNA c/ml |  |  | /mm<sup>3</sup> | HIV RNA c/ml |
| UL:<br>Ukwangiso Lwabantwana Study, South Africa | MTCT | 58 pairs | F | 23 yrs<br>(21-30) | 456<br>(255-566) | 8,000<br>(<20-770,000) | 42 female, 16 male | 37 hours<br>(21hrs-9d) | 1872<br>(1099-2446) | 18,000<br>(2,500-99,500) |
| PEHS: Paediatric Early HAART and Structured Treatment Interruption Study, South Africa | MTCT | 28 pairs | F | 27 yrs<br>(23-31) | 375<br>(211-483) | 49,750<br>(11,800-169,250) | 13 female, 15 male | 6 weeks<br>(4-8 weeks) | 1583<br>(1081-2108) | 91,550<br>(24,675-1,198,000) |
| FRESH: Females Rising through Education, Support, and Health, South Africa | ATAT | 15 recipients | N/A | N/A | N/A | N/A | 15 Female | 22 yrs<br>(21-23) | 642<br>(390-903) | 128,500<br>(6,250-380,000) |
| ZEHRP:<br>Zambia Emory HIV Research Project, Zambia | ATAT | 9 pairs | 4F, 5M | N/A | 500<br>(115-1008) | 79,264<br>(2,620-365,000) | 4M, 5F | N/A | 507<br>(320-828) | 2,941,142<br>(78,600-24,903,584) |

First, the authenticity of mother-child transmission pairs was validated by phylogenetic analysis of all the viral sequences in this study (Fig 1A). The VRC of chimeric viruses derived from the infants were then compared with those of the respective mothers. Overall, the VRC in the mothers was higher than in the children, both in the PEHSS cohort (mean VRC 0.85 versus 0.73; p=0.004) and in the UL cohort (mean VRC 0.91 versus 0.79; p=0.0003, Fig 1B).

**Figure 1:**
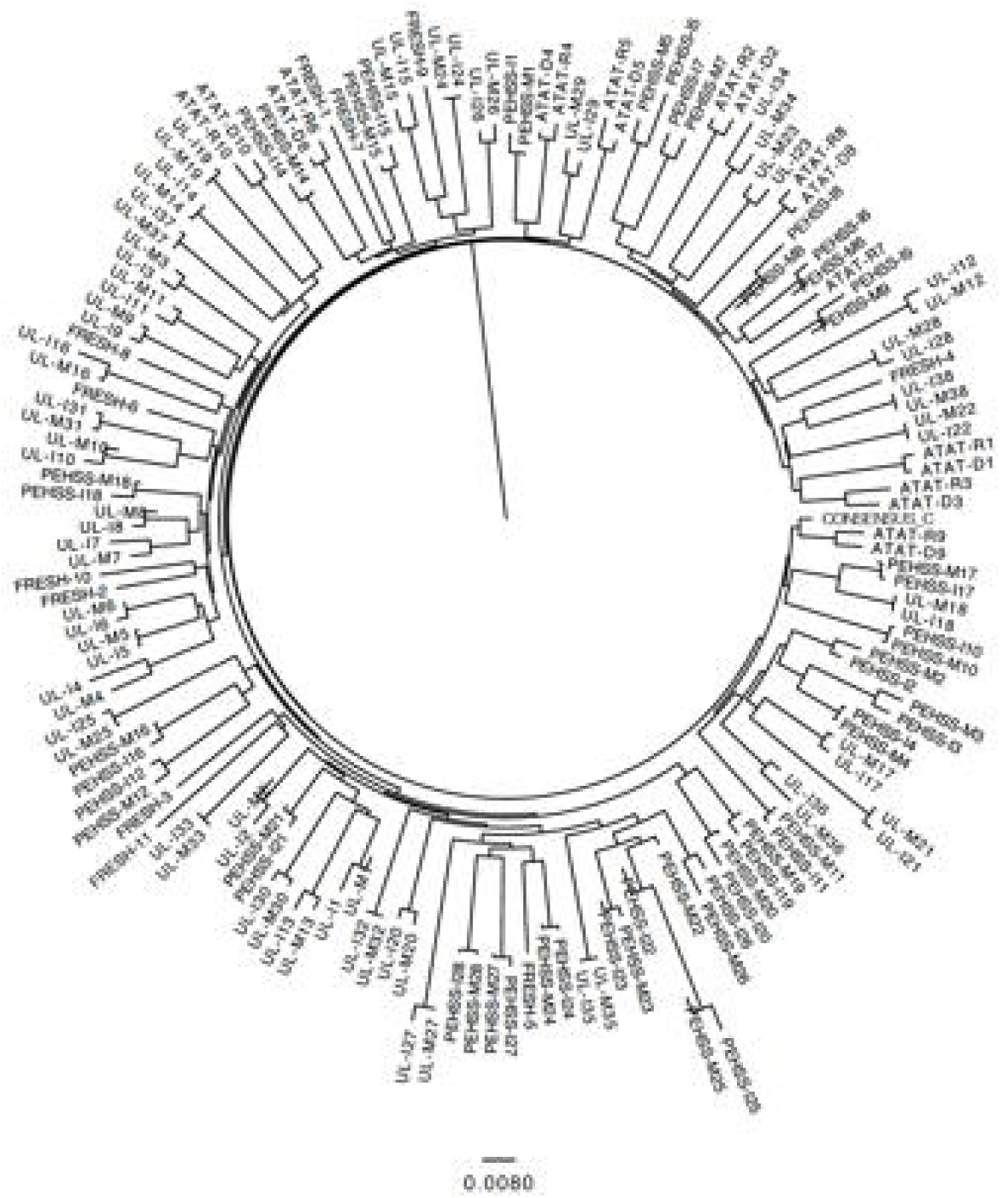
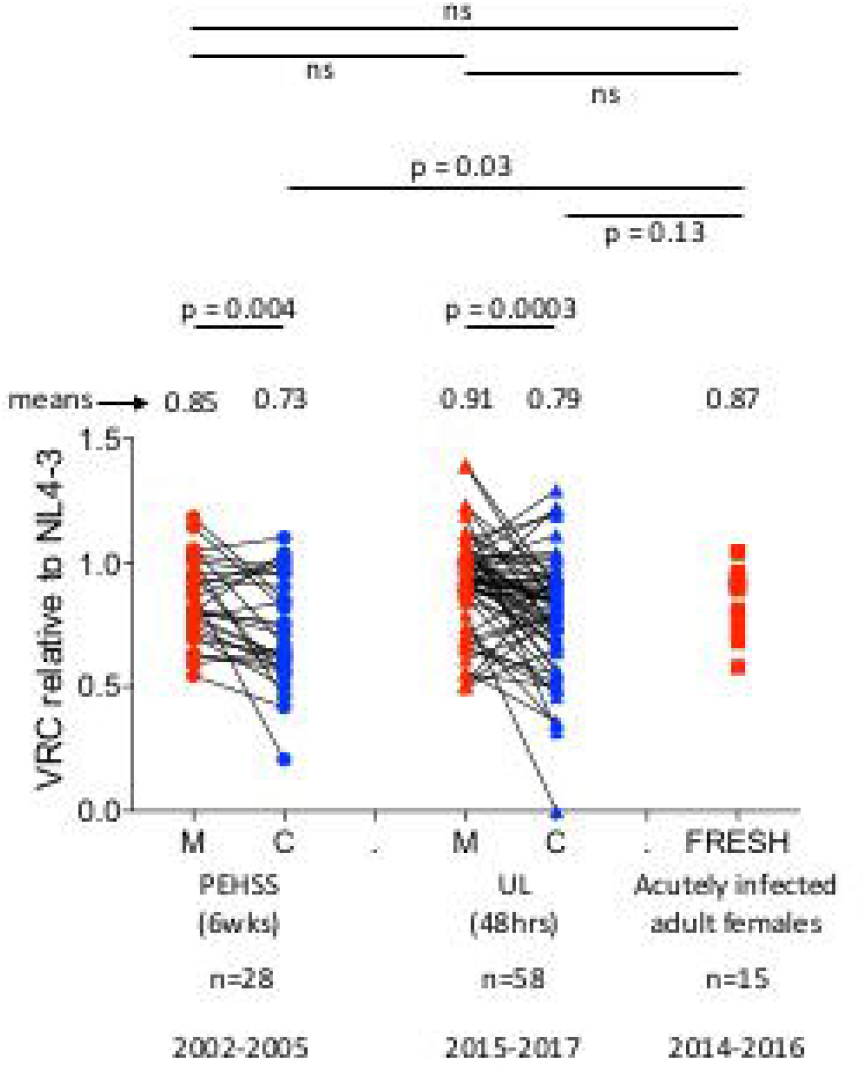
Transmission of low fitness viruses in MTCT of HIV-1. Replication capacities, normalized to NL4-3 comparator virus, were significantly lower in infants, mean 0.73, compared to mothers, mean 0.85 (n=28 pairs) in the PEHSS cohort (p=0.004), and in the UL cohort, infants mean 0.79, mothers mean 0.91 (n=58 pairs) (p=0.0003). Replication capacities in the acutely infected adult females (FRESH cohort), mean 0.87 (n=15) were not significantly different to the mothers in both the PEHSS and UL cohorts but were significantly higher than the infants of the PEHSS cohort (mean VRC 0.87 versus mean VRC 0.73, p=0.03).

The VRC of gag-protease chimeric viruses derived from the FRESH cohort acutely infected adult females had a mean of 0.87 (IQR 0.77-0.94, Fig 1B). These VRCs derived from the FRESH cohort did not differ significantly from those derived from the age-matched transmitting mothers in the PEHSS cohort (mean VRC 0.85, p=ns) or the UL cohort (VRC 0.91, p=ns, Fig 1B, Table 1), consistent with the notion that VRC in female HIV transmission donors and recipients do not differ significantly. However, the FRESH cohort VRCs tended to be higher than those of the MTCT recipient infants (p=0.03 and p=0.13, for FRESH versus the PEHSS cohort versus the UL cohort, respectively, Fig 1B).

### Differences in VRC between donor and recipient are greater in mother-to-child versus adult-to-adult heterosexual transmission of HIV

Recombinant viruses were generated from 9 heterosexual adult-to-adult transmission (ATAT) pairs from the Zambia Emory HIV Research Project (ZEHRP), Zambia, and gag-protease chimeric viruses were generated as described above. Five transmission pairs consisted of a male donor and female recipient and 4 transmission pairs comprised a female donor and male recipient. In contrast to what we observed in MTCT, there was no significant decrease in the VRC of the transmitted virus when compared to the donor VRC. The trend observed here in this small number of pairs studied favoured the transmission of a fitter virus (donor mean VRC 0.67 versus recipient mean VRC 0.72, p=ns, Fig 2A) in heterosexual ATAT. These are consistent with previous data generated from viral sequence analysis alone (22) indicating that fitter viruses are favoured more in woman-to-man than in man-to-woman transmission (male donor mean VCR 0.64 versus female recipient mean VRC 0.64, p=ns, Fig 2B; female donor mean VRC 0.70 versus male recipient mean VRC 0.83, p=ns, Fig 2C).

**Figure 2:**
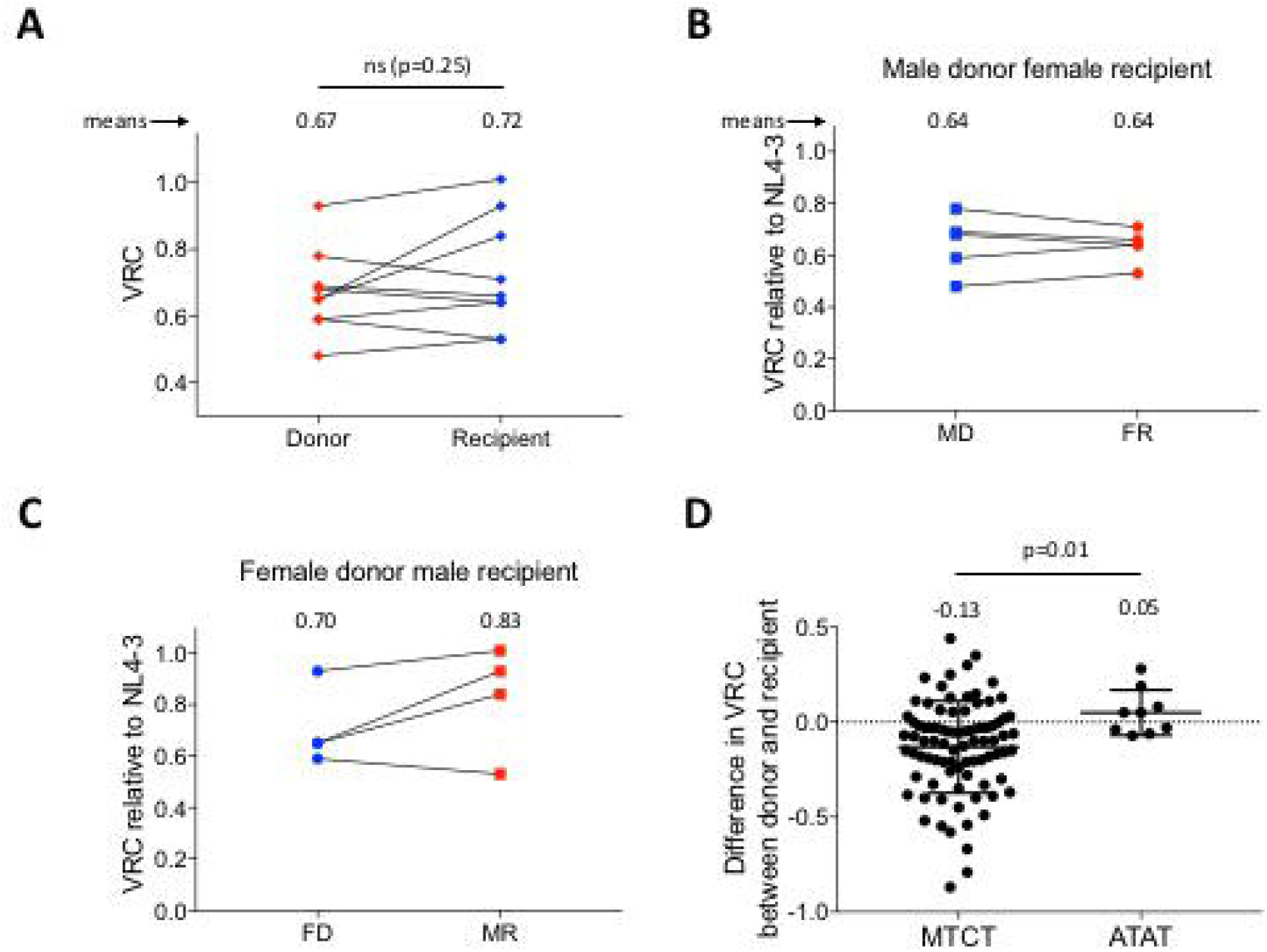
Preferential transmission of high fitness viruses in ATAT of HIV-1, particularly in female-to-male transmissions. [A] Replication capacities, normalized to NL4-3 comparator virus, in the ZEHRP acutely infected ATAT pairs (n=9) did not differ significantly between donor, mean VRC 0.67 and recipient, mean 0.72 (p=ns). [B] Male donor, female recipient ATAT pairs (n=5) did not differ in VRC, mean 0.64 versus 0.64 respectively. [C] Female donor, male recipient ATAT pairs (n=4) favoured transmission of fit viruses, mean VRC 0.70 versus mean VRC 0.83, respectively. [D] Difference in VRC between donor and recipient in MTCT (n=86) versus ATAT (n=9) (p=0.01)

We next compared the differences in VRC between donor and recipient isolates among the ATAT pairs with MTCT pairs. Here we observed that, despite the small number of adult transmission pairs studied, there was significant difference (p=0.01, Fig 2D), with the transmitted viruses in ATAT pairs similar to, or somewhat higher than, donors’ viruses, whereas in MTCT the transmitted viruses are typically of lower VRC when compared to those of the donors.

### Single genome amplification (SGA) of gag-protease chimeric viruses corroborates preferential transmission of lower replicative capacity viruses in MTCT

A possible explanation for this assay to provide an artefactually inflated estimate of the donor VRC might be that the donor viral quasispecies likely exhibit a degree of diversity, and the chimeric viruses generated by co-transfection of the amplified patient-derived gag-protease genes with linearized pNL4-3Δgag-protease could potentially be biased in favour of the faster growing, highest VRC viruses. By contrast, the recipient viral quasispecies would typically have a much lower diversity, and hence would not be so prone to bias in favour of rapidly growing, high VRC isolates. The outcome from this potential bias would be an apparent drop in VRC when comparing donor with recipient virus, as we have described here and previously in MTCT.

To address this issue, we performed limiting dilutions of patient RNA to obtain a diverse repertoire of clonal gag-protease isolates. Single genomes were confirmed by sequencing and transfected with linearized pNL4-3Δgag-protease into GFP reporter cell lines as described above. We first generated clones via single genome amplification (SGA) from 8 mother-infant pairs from the UL cohort. Among the 8 mothers, the maternal gag-protease population-generated VRC strongly correlated with the SGA-derived VRC (r=0.973, p=<0.0001, Fig 3AB) and a similar correlation was observed between the VRCs generated by the infant gag-protease population and the VRC determined from the clonal viral isolates (r=0.972, p=<0.0001, Fig 3CD). The Gag-Pro population-derived VRCs in both cases were indeed higher than Gag-Pro SGA-derived VRCs (mean difference 0.045 in the mothers and 0.027 in the infants), however in neither case did these reach statistical significance, nor did they explain the magnitude of VRC differences (mean 0.13) observed between mother and infant VRC (Fig 2D).

**Figure 3:**
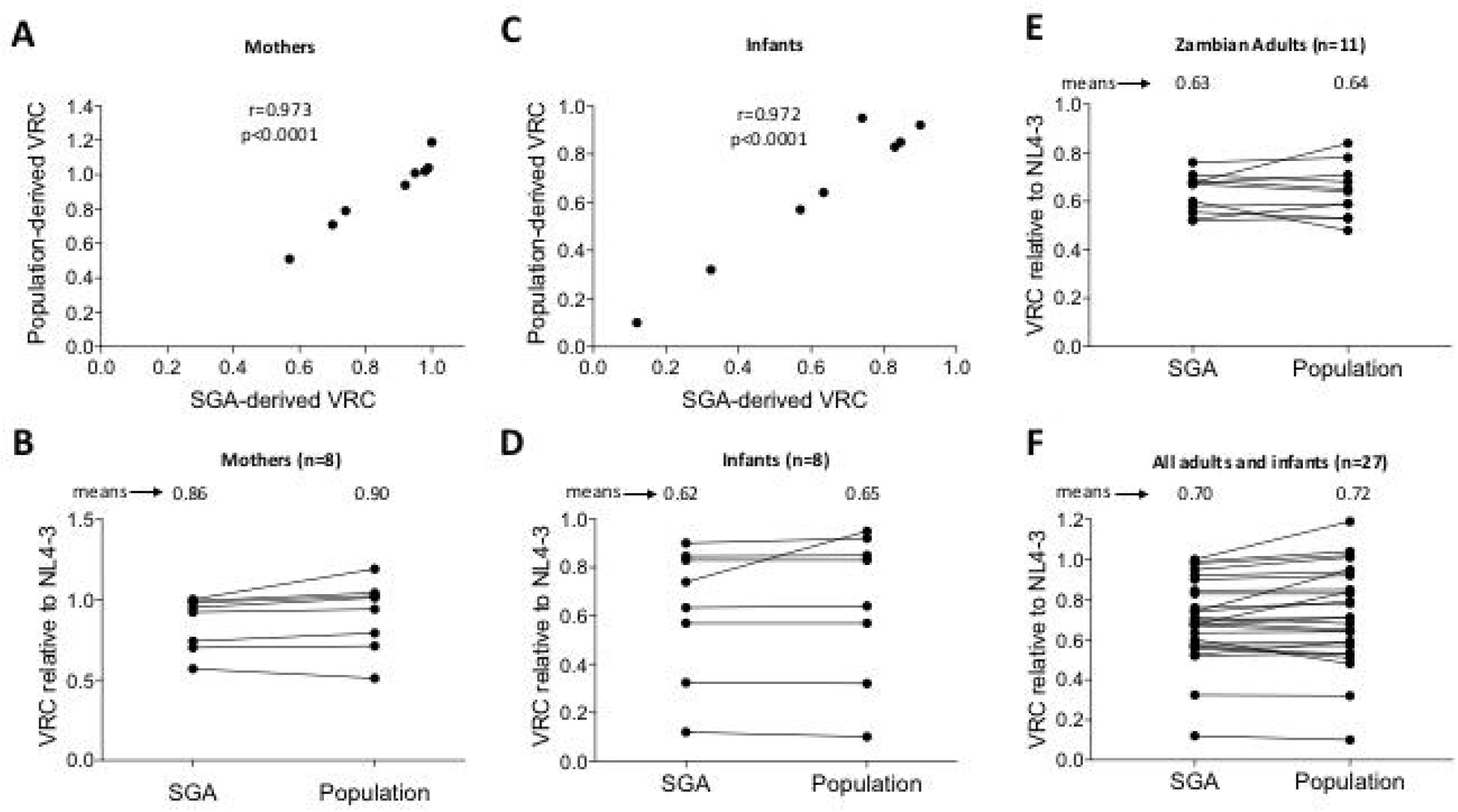
SGA of viral isolates corroborates a selection bias of low fitness viruses in MTCT and high fitness viruses in ATAT HIV transmission. Spearman correlations between population VRC and the mean VRC obtained from single viral isolates in [A] UL mothers (n=8) (r=0.973, p=<0.0001) and [B] UL infants (n=8) (r=0.972, p=<0.0001). Paired t test of population VRC and the mean VRC obtained from single viral isolates in [C] UL mothers (n=8) (mean difference 0.045, p=0.11) and [D] UL infants (n=8) (mean difference 0.027, p=0.34). Paired t test between population VRC and the mean VRC obtained from single viral isolates in [E] ZEHRP adults, mean 0.64 versus 0.63 respectively (n=10) (mean of differences 0.005, p=ns) and [F] ZEHRP ATAT pairs and UL MTCT pairs (n=26), mean 0.72 versus 0.70 (mean of differences 0.02, p=ns).

To further test the hypothesis that the drop in VRC between donor and recipient observed in MTCT can be explained by the assay, we conducted the same evaluation comparing Gag-Pro population-derived VRCs with Gag-Pro SGA-derived VRCs in 11 individuals within the 9 heterosexual ATAT pairs studied from the ZEHRP cohort. Importantly, as with all the VRC measurements, these were conducted blind to study subject identification. Among 5 donors analysed, the Gag-Pro population-derived VRCs was marginally lower than the Gag-Pro population derived VRCs (mean −0.012), and among the 6 recipients, the Gag-Pro population-derived VRCs was marginally higher than the Gag-Pro population derived VRCs (mean +0.02). Overall, the mean difference in these 11 adults between Gag-Pro population-derived VRCs and the Gag-Pro SGA-derived VRCs was only 0.005 (Fig 3E). Combining all of these comparisons for 27 individuals evaluated, from whom a median of 6 clones per subject were evaluated (IQR 5-7), on average the Gag-Pro population-derived VRCs were marginally and not statistically significantly higher than SGA-derived VRCs (mean VRCs 0.72 versus 0.70, respectively, ie a VRC difference of 0.02). These findings would indicate, first, that the drop in VRC observed in MTCT is indeed real and not a consequence of the assay used; and, second, that the difference between donor and recipient VRC in MTCT compared with ATAT is also non-artefactual and statistically significant.

## DISCUSSION

These analyses of the replication capacity of viruses transmitted from MTCT, compared with those in ATAT, were designed to address the question of whether the reported reduction in VRC of transmitted virus in MTCT represents a significant difference from what is observed in ATAT. These comparisons have not been done previously. Although the studies that have been undertaken would suggest that fundamental differences do exist in respect of the viruses transmitted through the respective bottlenecks, a question arises over whether the Gag-Pro-driven assay, that has been used in MTCT studies to measure viral replicative capacity (VRC), might artefactually inflate the donor VRC, and hence also the difference between donor and recipient virus. The rationale behind this hypothesis is that the viruses in the donor would typically be more diverse in VRC than those transmitted to the recipient, and hence the culture of Gag-Pro-NL4-3 chimeric viruses might be expected to favour fast-growing, high VRC viruses within this donor pool. By contrast, this step of viral culture might have less impact among recipient viruses comprised largely of very similar transmitted/founder-like viruses.

If this effect were to have a significant bearing on the measurement of donor VRC, this would artefactually increase both donor VRC and the difference between donor and recipient VRC. In this way, the reported lower VRC of MTCT virus in the child compared to the mother (7, 8) might in reality not exist. To address this possibility, we here extended the original findings that were originally described in South African mother-child cohorts comprising children aged 7.5yrs (7) and infants aged 6 weeks (8), to another cohort of mother-child pairs from KwaZulu-Natal, South Africa, in which the *in utero* infected child was aged 36 hours (median) at the time the sample was taken. The findings in the newborns show a result very similar to that previously observed in the South African infants aged 6 weeks, consistent with the conclusion that these differences in VRC between mother and child arise as a consequence of the MTCT bottleneck, and not subsequent to transmission.

To determine whether the VRC of viruses in the transmitting mothers was higher than in the acutely infected, age-matched recipient females, we had included in these analyses samples from 15 females (median age 22yrs) in the FRESH cohort, also from KwaZulu-Natal South Africa, all but one of whom were enrolled in Fiebig Stage I (15, 23). The VRC of the FRESH cohort viruses was not significantly different from that of the mothers in the two mother-child cohorts studied, but did differ from VRC in the infants (p=0.03 in the PEHSS cohort and p=0.13 in the UL cohort). Although the VRC of the donors in the FRESH cohort was unknown, these data were consistent with the VRC of transmitted virus in MTCT being lower than in the donor, but in ATAT being similar to that in the donor.

To evaluate this directly in heterosexual ATAT pairs, we studied 9 pairs within the ZEHRP Zambian cohort (2–4, 9, 24). As in the mother-child pair studies, VRCs were determined blind to the identity of the study subjects. Following unblinding, we showed that the VRCs of virus transmitted in heterosexual ATAT are similar to, or slightly higher than, viruses in the donor. This result is therefore consistent with previous studies using different methodologies to measure VRC (9, 10), and, despite the small numbers of adult pairs evaluated, a statistically significant difference between VRC in donor and recipient virus was observed between the MTCT and the heterosexual ATAT pairs. These data indicate that any artefactual inflation in the donor VRC resulting from the Gag-Pro assay adopted is too small to obscure a fundamental difference in the MTCT and ATAT bottleneck.

To address the question of whether Gag-Pro population-generated VRC are significantly higher than Gag-Pro SGA clone-derived VRC, we undertook these analyses in donor and recipient of the MTCT and ATAT pairs and showed that there is a mean difference of 0.02 in VRC across 27 subjects evaluated (populated-derived VRC being 0.02 higher). This difference was not statistically significant and is too small to explain the 0.13 difference in VRC between donor and recipient virus in MTCT.

Although the numbers in the Zambian cohort were too small to evaluate potential sex differences in the VRC of adult-to-adult transmitted viruses, there is a hint from these preliminary data that females are more permissive to infection from lower VRC viruses, whereas the viruses infecting males are typically higher fitness viruses. If true, these data would be consistent with the conclusions based on analysis of viral sequences alone, that did not measure VRC directly but estimated viral fitness from the frequency of a variant in the same Zambian study cohort (25). Further studies to evaluate these potential sex differences in heterosexual ATAT and the mechanisms underlying this apparent increased susceptibility to infection among females would be important, particularly as epidemiological studies have consistently argued that the risk of heterosexual HIV transmission from female-to-male is the same as from male-to-females, after confounding factors such as viral load and the presence of sexually transmitted diseases have been taken into account (25–29).

These studies now confirm previous findings that the virus transmitted to the infant *in utero* has a substantially lower VRC than that of virus in the mother. This is an important result, especially in considerations of HIV cure, since transmission of viruses with low VRC has been shown to result in lower levels of immune activation and smaller viral reservoirs (3). Also, it has been hypothesised that *in utero* infected children may have greater potential for post-treatment control or even true eradication of the virus (30–32). Two of the main reasons for this suggestion are that, first, *in utero* infected children can be treated with antiretroviral therapy within hours of birth and therefore very early after infection; and, second, the tolerogenic immune environment of early life does not ‘fuel the fire’ of acute infection in the way that the highly activated adult immune response does (5, 6, 33, 34). Both of these factors would tend to reduce the size and diversity of the initial viral reservoir. To these may now be added a third factor, that of a reservoir comprised of low VRC virus, that also would be expected to contribute to increased cure potential among *in utero* infected infants (3).

Precisely what factors contribute to the differential transmission bottleneck in MTCT and ATAT remain largely unknown and in MTCT have been relatively unstudied. Whilst current interventions have reduced the number of new infections by almost half (to 1.8m/year: http://www.unaids.org/en/resources/fact-sheet) since their peak in 1996, defining these mechanisms remain fundamental to the development of new and effective prevention strategies to reduce the numbers of new infections further.

## ACKNOWLEDGEMENTS

Funding for this work came from grants to PG from the Wellcome Trust (Grant 104748MA) and the NIH (RO1 AI133673).

